# Association of EBV (Type 1 and 2) with Histopathological Outcomes in Breast Cancer in Pakistani Women

**DOI:** 10.1101/2021.11.16.468790

**Authors:** Yusra Ilyas, Sanaullah Khan, Naveed Khan

## Abstract

**Introduction:** Breast cancer is one of the major and frequent tumors in the public health sector globally. The rising global prevalence of breast cancer has aroused attention in a viral etiology. Other than genetic and hormonal roles, viruses like Epstein - Barr virus (EBV) also participate in the development and advancement of breast cancer.

**Aim:** This study was conducted to detect the frequency of EBV genotypes in breast cancer patients and compare it with histopathological breast cancer changes.

**Methods:** Formalin-fixed paraffin-embedded samples of breast cancer (N=60) ages ranged from 22-70 years were collected. EBV DNA was isolated, amplified, typed through PCR, and correlated with histopathological outcomes of breast cancer using SPSS software version 26.

**Results:** Our findings suggest that among breast cancer factors, Invasive ductal carcinoma (IDC) was the most common pathological pattern found among patients (90%), observed statistically significant (p= 0.01275). In regards to clinical staging, 8 (13.3 %) patients diagnosed with stage I, 39 (65 %) with stage II, and 13 (21.6 %) with stage III reported statistically significant association (p=0.0003). EBV DNA was detected in 68.3% (41/60) breast cancer patients, reported a statistically significant difference between the prevalence of EBV in breast cancer patients and normal samples (*p* = 0.001). Of 41 EBV-positive samples, 40 were EBV-1, while only 1 had EBV-2 infection (p < 0.001). No influence on cancer histology was observed. Regarding the association of breast cancer with EBV, histological type (P =0.209), tumor stage (P = 0.48), tumor grade (0.356), tumor sizes (p= 0.976), age (p= 0.1055), tumor laterality (p= 0.533) and ER/PR status (p=0.773) showed no significant association.

**Conclusion:** EBV-1 is prevalent in breast cancer patients and associated with IDC in the study area. For conclusive evidence, more studies are required based on a large sample size and by using more sensitive techniques.

## 1. Introduction

Breast cancer is globally accounted for 23% of all cancer cases (Asif et al., 2014), currently representing significant public health issues worldwide. It is considered metastatic cancer that can also be spread to far-flung organs, including bones, liver, lungs, and the brain (Ahmad, 2019). Breast cancer’s molecular characteristics include human epidermal growth factor receptor 2 (HER2) activation, hormone receptor activation (estrogen receptor or ER and progesterone receptor or PR), and mutations in BRCA. About 800,000 incidences have been annually reported affecting them worldwide, thus making it the most common neoplastic disease of women (McPherson *et al*., 2000). Several internal and external risk factors may raise the likelihood of developing breast cancer, including genetic predisposition (e.g., mutations in BRCA1/2 and other genes), a family history of breast cancer, ethnicity (more common in the Caucasian population), dense breast tissue, lifestyle, obesity (Alibek et al., 2013), insulin resistance, and to reproductive background including age at first gestation, early menarche, late menopause, hormonal therapy, null parity and aging (Schwingel et al., 2019), however, identification of the exact causes is complex. Viral infections are considered biological carcinogens are contributing about 18-20 percent of all cancers related to humans (Hippocrate *et al*., 2011; Alibek et al., 2013). Several recent studies reported viruses like Epstein-Barr Virus, papillomavirus, mouse mammary tumor virus, bovine leukemia virus (Lawson and Heng, 2010; Fozuni et al., 2020), and Human Herpes Virus-8 (Fozuni et al., 2020) had been implicated in the development of breast cancers. All these have known oncogenic potential and have been detected in normal and malignant breast tissues (Lawson and Heng, 2010).

Epstein-Barr Virus (EBV), also known as human gammaherpesvirus 4 (Jean-Pierre et al., 2012), is included among the eight known human herpesviruses and is classified as a class I carcinogen by IARC (Alibek et al., 2013), as it is associated with the pathogenesis of breast cancer (Glaser et al., 2017; Dumitrescu *et al*., 2005; Fessahaye et al., 2017). EBV is considered an element of danger for the abundant incidences of breast cancer as the statistical data exhibits its resemblance with Hodgkin’s disease in young adulthood (Hippocrate *et al*., 2011). EBV is estimated to be responsible for 1.5% of all cancer worldwide, though virus persistence is often asymptomatic and does not develop any disease (Jean-Pierre et al., 2012). Gulley (2001) reported that most asymptomatic carriers harbor up to 50 EBV genomes per million B cells.

EBV primarily infects epithelial, B-cells, and human mammary epithelial cells (MECs) (Hu *et al*., 2016). EBV encompasses a 172 kbp DNA genome with a linear double-stranded structure that encodes more than 85 genes (Böhm, 2010). It is lymphotropic in nature and is a principal cause of infectious mononucleosis (Johannsen *et al*., 2015). EBV has two significant genotypes, types 1 and 2, differentiated based on the EBNA gene. The EBNA gene is held responsible for gene regulation and viral DNA replication (Smatti *et al*., 2018; Farrell *et al*., 2019). One of the clinical differences is that EBV 1 can convert B cells into lymphoblastoid cell lines (LCL) more efficiently than EBV 2 (Zanella et al., 2019). From the discovery of EBV in 1964 and to understanding its etiological in causing infectious mononucleosis in 1968, large-scale epidemiological studies have been conducted globally to investigate the epidemiology of EBV (Sharifipour and Rad, 2020). It has been shown that few EBV oncoproteins are expressed in different types of cancer, i.e., EBNA1 and EBBR are expressed in Burkett’s lymphoma, EBNA1, LMP1, LMP2, EBERs expressed in nasopharyngeal carcinoma, EBNA 1, LMP2, EBERs expressed in gastric Carcinoma (Li *et al*., 2010). Epstein Barr-Nuclear antigen EBNA-3C is displayed at the latency stage and may turn primary B lymphocytes into lymphoblastic cell lines. EBNA 3C targets many tumor suppressors such as p53, p73, and pRB via a phosphorylation and ubiquitination pathway in the course of transformation (Sharifpour *et al*., 2019).

The relationship of several human lymphomas, such as Hodgkin and Burkitt’s lymphoma, nasopharyngeal carcinomas, certain epithelial cell carcinomas, etc., with EBV, has been revealed by various modern studies (Salahuddin *et al*., 2018; Fawzy *et al*., 2008). Breast epithelial cells can be infected by EBV through direct contact with EBV-bearing lymphoblastoid cells (Hu *et al*., 2016). Different reports have been submitted concerning the presence of viruses in breast tissue, which differs greatly concerning the nature of the pre-analytical process (e.g., type of sample, type of storage, sample type, type of storage, type of sample) (Gannon *et al*. 2018). The TRADD (TNF-receptor 1-associated death domain protein) gene is an essential factor for the function of the EBV virus-encoded latent membrane protein 1(LMP1), causing uncontrolled proliferation of EBV infected cells and ultimately cancer (Lawson and Heng, 2010).

The risk of breast cancer is growing in developing countries, where 44% of the world’s breast cancer deaths and 39% of overall new breast cancer cases are diagnosed in Asia (Madhav et al., 2018). In Asia, Pakistan has a high burden of breast cancer (Menhas and Umer, 2015; Majeed et al., 2014; Zaheer et al., 2019) among Asian countries (Bano *et al*., 2016), whereby one in every nine women has a lifetime risk of being diagnosed with breast cancer (Asif et al., 2014; Zaheer et al., 2019). Pakistan, as a developing country, faces many challenges regarding awareness, screening, late presentation, and its management, especially in rural areas of Pakistan, where lower education, literacy, and low socioeconomic conditions are responsible for the poor health of women. Mammograms are an effective but costly screening program that is unaffordable for most of the population, leading to the presentation of breast cancer later (Menhas and Umer, 2015). In Pakistan, one in every nine women suffers from breast cancer, 2.5 times higher than India and Iran’s neighboring countries (Menhas and Umer, 2015; Majeed et al., 2014). A study on the incidence and mortality of breast cancer and their relationship to development in Asia reported that the highest age-standardized death rate (2.52) due to breast cancer was observed in Pakistan (Ghoncheh et al., 2015). A relationship between breast cancer and EBV is potentially important for understanding breast cancer etiology, breast cancer treatment, early detection, and prevention (Glaser et al., 2004). EBV-1 has been reported in various cancer patients in Pakistan (Salahuddin *et al*., 2018). Still, only one study has identified EBV in breast cancer patients (Naushad *et al*., 2017), but EBV genotypes have not been identified in connection to breast cancer. Moreover, the correlation of histopathological changes with EBV genotypes has not been reviewed yet. Therefore, this study was aimed to determine if there is a higher association between breast cancer and the presence of EBV genotypes in breast cancer patients of Pakistan and its comparison with histopathological outcomes of Breast cancer.

## 2. Materials and methods

### 2.1. Study Subjects and Questionnaire Content

Formalin-fixed paraffin-embedded (FFPE) samples were collected from 60 patients from various hospitals of Peshawar, Pakistan, where samples of breast cancer patients from different regions of Khyber Pakhtunkhwa were diagnosed during 2019-2020.

The diagnosed breast cancer patients (60 cases) were selected as the breast cancer group while the sample of 20 healthy donors was collected to categorize as a control group following the 1:3 principles, taking age as a matching factor. Members of the control group were selected after breast examination by a skillful physician. For baseline data collection, a questionnaire was designed having all the primary information. Clinical information (cancer stage, grade of tumor, histological subtype, tumor laterality, estrogen, and progesterone receptors status) was retrieved from the clinical file.

### 2.2. Histopathological Information

Histopathological analysis of microscopic slides and clinical information was concluded by a consultant histopathologist. They examined the tumor grading based on cellular differentiation and resemblance to its standard counterpart and staging based on the tumor’s extent and spread involvement. Histopathological features were documented according to a standardized format using microscopic observations based on invasive cancer size, grade based on the degree of differentiation (i.e., nuclear pleomorphism), histological subtype according to the growth pattern of tumor, cellular pleomorphism, and proliferation (i.e., mitotic index).

### 2.3. DNA extraction and amplification

DNA from the deparaffinized tissues was isolated through the xylene ethanol method and stored at −70 °C by using a high pure PCR template preparation kit (Thermofisher, USA) according to the manufacturer’s instructions. The DNA extraction from samples was done by the earlier mentioned protocol of Sharifpour et al. 2019. The synthesized cDNA was amplified by EBNA 3C (1F and 1R) primers (5’-GAGAAGGGGAGC GTGTGTTGT-3’, 5’-GCTCGTTTTTGACGTCGGC-3’) by using regular PCR and adding *Taq* DNA polymerase (ThermoFisher, USA). The components of 25 μL reaction mixture contained 10 μL extracted DNA, 2.5μL PCR buffer 10X, 3 μL MgCl2 25mM, 0.5μl dNTP 10 mM (Roche), 6.8 μL dH_2_O,0.2μL Taq Polymerase (Roche),1 μL of each primer sequence which were added to thermocycler under the conditions of initial denaturation at 95 °C for 10 min, denaturation, annealing and extension at 94 °C for 45 sec, 55 °C for 45 sec, 72 °C for 45 sec (35 cycles) and final extension at 72 °C for 10 min. By nested PCR, the synthesized cDNA was re-amplified by EBNA 3C (2-F and 2-R) primers (Table-1). The second round was carried out with 4μl of the first-round product, under the same condition described previously with the set of different nested PCR primers.

**Table 1.**
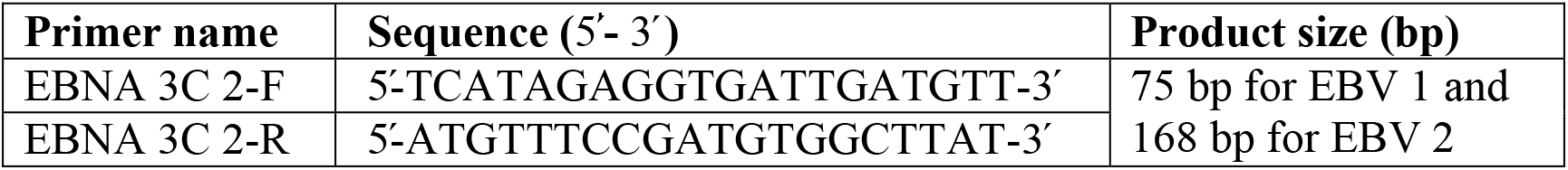
Primer sequence for cDNA formation

### 2.4. Electrophoresis

In 0.5 ×TBE (Tris Borate EDTA buffer), 2% agarose gel was prepared by adding 3 μL ethidium bromide and then run-on nested PCR product. To each of the nested PCR products, 2 μL loading dye was applied. 50 bp DNA ladder marker (5 μL) (Fermentas, USA) was also added parallel to samples in each row and were run for 45 minutes (90 volts, 500 Amperes). Nested PCR products were 75 bp for type 1 and 168 bp for type 2 EBV. cDNA bands under UV transilluminators were compared and visualized with DNA ladder markers.

### 2.5. Data analysis

All patients were categorized based on age, breast cancer type, tumor grade, tumor size, tumor laterality, and ER/PR status. The study population was divided into three age groups; 22-39 years, 40-59 years, and ≥60 years. On the basis of histology, breast cancer was divided into four types, Invasive ductal carcinoma (IDC), Invasive lobular carcinoma (ILC), metaplastic carcinoma, and secretory carcinoma. The study population was classified on the basis of tumor size into three groups: ≥2.9cm, 3-5.9cm, and ≥6cm and according to the TNM staging of the American Joint Committee on Cancer, breast cancer is divided into three tumors grades: I, II, III. The prevalence of EBV related to these specific factors was calculated, indicating the role of EBV in breast cancer progression.

### 2.6. Statistical analysis

The obtained results were analyzed through SPSS software version 26. Chi-square (χ^2^) test or Fisher’s exact test, where appropriate, was used for analysis between EBV and clinic-pathological data and odds ratio (OR) with a 95% confidence interval (CI). Statistical significance was achieved at *p < .05*.

## 3. Results

### 3.1. Clinicopathological characteristics of patients with breast cancer

The breast tissues were collected from all patients and the control group and analyzed for EBV prevalence and genotypes. The age range and mean (±SD) age of the breast cancer patients was 22-70 years and 47.783± 12.082 years, and the control group was 22-70 years and 58.92 (18±19) years; showed no significant difference between the two groups (p=0.938). Clinicopathological characteristics of patients with breast cancer were presented in table-2. The patients in the age group of 40-59 years had the highest rate of breast cancer followed by 22-39 years and ≥ 60 years with the rate of 48.3%, 26.6%, and 25%, respectively, showed statistically significant association (p=0.001). Regarding the histological type, IDC was the most common pathological pattern found among patients (90%), followed by ILC (n=3, 5%), metaplastic carcinoma (n=2, 3.33%), and secretory carcinoma (n=1, 1.66%), observed a statistically significant association between age and types of breast cancer disease (p= 0.01275). Regarding the clinical staging, 8 (13.3 %) patients diagnosed with stage I, 39 (65 %) with stage II, and 13 (21.6 %) with stage III reported statistically significant association (p=0.0003). The mean age of patients of grade I, grade II, and grade III was 60.25 (± 8.59), 50.69 (±12.70), and 40 (± 8.26) years old, respectively, and reported significant association (p=0.011). The size of the tumor of patients ranged from 0.1 to 8.5cm. Fifty percent (30) patients have tumor size ≥ 2.9cm, 38.3% (23) have tumor size 3-5.9cm while 7 (11.6 %) have ≥ 6cm. The largest and smallest tumor was reported in grade III (37 years old) and grade II (35 years old) patients positive of IDC, respectively. The tumor grade with different age groups showed significant association (p=0.095). Of these, 55% (n=33) of patients showed involvement of right breast, while 45% (n=27) cases showed left breast involvement, showed significant association (p=0.011). Our data revealed that age (p=0.001), tumor stage (p=0.0003) and tumor size (p=0.02) showed significant association while type of breast cancer disease (p=0.1), tumor laterality (p=0.9) and ER/PR status (p=0.6) showed no significant association. Statistically, age groups and tumor grade of breast cancer patients (p=0.005) and age groups, tumor grade, and tumor laterality showed significant association (p=0.005). Table 3 showed the statistical association of Tumor Grade, Age Group, ER/PR Status, and tumor laterality.

**Table 2.** Clinic pathological characteristics of patients with breast cancer.

| Characteristics | Value N (%) | p-value |
| --- | --- | --- |
| Age group (years) |  |  |
| I | 16 (26.6) | 0.001 |
| II | 29 (48.3) |  |
| III | 15 (25) |  |
| Type of BC disease |  |  |
| IDC | 54 (90) | 0.01275 |
| ILC | 3 (5) |  |
| Metaplastic carcinoma | 2(3.33) |  |
| Secretory carcinoma | 1(1.66) |  |
| Tumor grade |  |  |
| I | 8 (13.3) | 0.0003 |
| II | 39 (65) |  |
| III | 13 (21.6) |  |
| Tumor size (cm) |  |  |
| 0-2.9 | 30 (50) | 0.02 |
| 3- 5.9 | 23 (38.3) |  |
| > 6 | 7 (11.6) |  |
| Tumor laterality |  |  |
| Right | 27 (45) | 0.011 |
| Left | 33 (55) |  |
| ER/ PR status |  |  |
| Positive | 29 (48.3) | 0.6 |
| Negative | 31 (51.6) |  |

**Table 3.** Statistic association of Tumor Grade, Age Group and ER/PR Status and tumor laterality

| Association of Tumor Grade, Age Group, and ER/PR Status |  |  |  |  |  |  |  |
| --- | --- | --- | --- | --- | --- | --- | --- |
| ER/PR Status |  |  | Tumor Grade |  |  | Total | p-value |
|  |  |  | I | II | III |  |  |
| Positive | Age Group | 22-39 | 0 | 4 | 3 | 7 | 0.330 |
|  |  | 40-59 | 2 | 10 | 3 | 15 |  |
|  |  | ≥60 | 0 | 8 | 1 | 9 |  |
|  | Total |  | 2 | 22 | 7 | 31 |  |
| Negative | Age Group | 22-39 | 1 | 4 | 4 | 9 | 0.052 |
|  |  | 40-59 | 5 | 7 | 2 | 14 |  |
|  |  | ≥60 | 0 | 6 | 0 | 6 |  |
|  | Total |  | 6 | 17 | 6 | 29 |  |
| Total | Age Group | 22-39 | 1 | 8 | 7 | 16 | 0.011 |
|  |  | 40-59 | 7 | 17 | 5 | 29 |  |
|  |  | ≥60 | 0 | 14 | 1 | 15 |  |
|  | Total |  | 8 | 39 | 13 | 60 |  |
| Association of tumor laterality, tumor grade, and age groups |  |  |  |  |  |  |  |
|  |  |  | Tumor Grade |  |  |  | p-value |
| Tumor Laterality |  |  | I | II | III | Total |  |
| Right | Age Group | 22-39 | 1 | 2 | 6 | 9 | 0.005 |
|  |  | 40-59 | 5 | 9 | 3 | 17 |  |
|  |  | ≥60 | 0 | 7 | 0 | 7 |  |
|  | Total |  | 6 | 18 | 9 | 33 |  |
| Left | Age Group | 22-39 | 0 | 6 | 1 | 7 | 0.574 |
|  |  | 40-59 | 2 | 8 | 2 | 12 |  |
|  |  | ≥60 | 0 | 7 | 1 | 8 |  |
|  | Total |  | 2 | 21 | 4 | 27 |  |
| Total | Age Group | 22-39 | 1 | 8 | 7 | 16 | 0.011 |
|  |  | 40-59 | 7 | 17 | 5 | 29 |  |
|  |  | ≥60 | 0 | 14 | 1 | 15 |  |
|  | Total |  | 8 | 39 | 13 | 60 |  |
| Association of ER/PR Status, Tumor Grade, and Tumor Laterality |  |  |  |  |  |  |  |
| ER/PR Status |  |  | Tumor Grade |  |  |  |  |
|  |  |  | I | II | III | Total | p-value |
| Positive | Tumor Laterality | Right | 1 | 10 | 1 | 12 | 0.464 |
|  |  | Left | 5 | 10 | 4 | 19 |  |
|  |  | Total | 6 | 20 | 5 | 31 |  |
| Negative | Tumor Laterality | Right | 1 | 11 | 3 | 15 | 0.500 |
|  |  | Left | 1 | 9 | 4 | 14 |  |
|  |  | Total | 2 | 20 | 7 | 29 |  |
| Total | Tumor Laterality | Right | 2 | 21 | 4 | 27 | 0.424 |
|  |  | Left | 6 | 19 | 8 | 33 |  |
|  |  | Total | 8 | 40 | 12 | 60 |  |

### 3.2. Histopathological Analysis of Breast Cancer Images

Performing the histopathology analysis of images, the features of keen interest to pathologist refers to cells size and nuclei shape, their spatial arrangement and organization into tubules, interaction with the stroma, etc. Fig 2a showed the normal breast tissue slide image with proper ductal and lobular system surrounded by fatty cells and connective tissue stroma. Whereas Fig 2b and 2c illustrated IDC images showing a disrupted pattern of growth and cellular morphology. Sections revealed breast tissue showing infiltration by a malignant neoplasm. The neoplasm is composed of the duct and tubule-like structures lined by atypical cells having hyperchromatic pleomorphic nuclei, prominent nucleoli, and a high N/C ratio. Also, no evidence of necrosis was observed. No such cellular difference was observed apparently among the images of EBV positive and negative samples.

**Fig 1.**
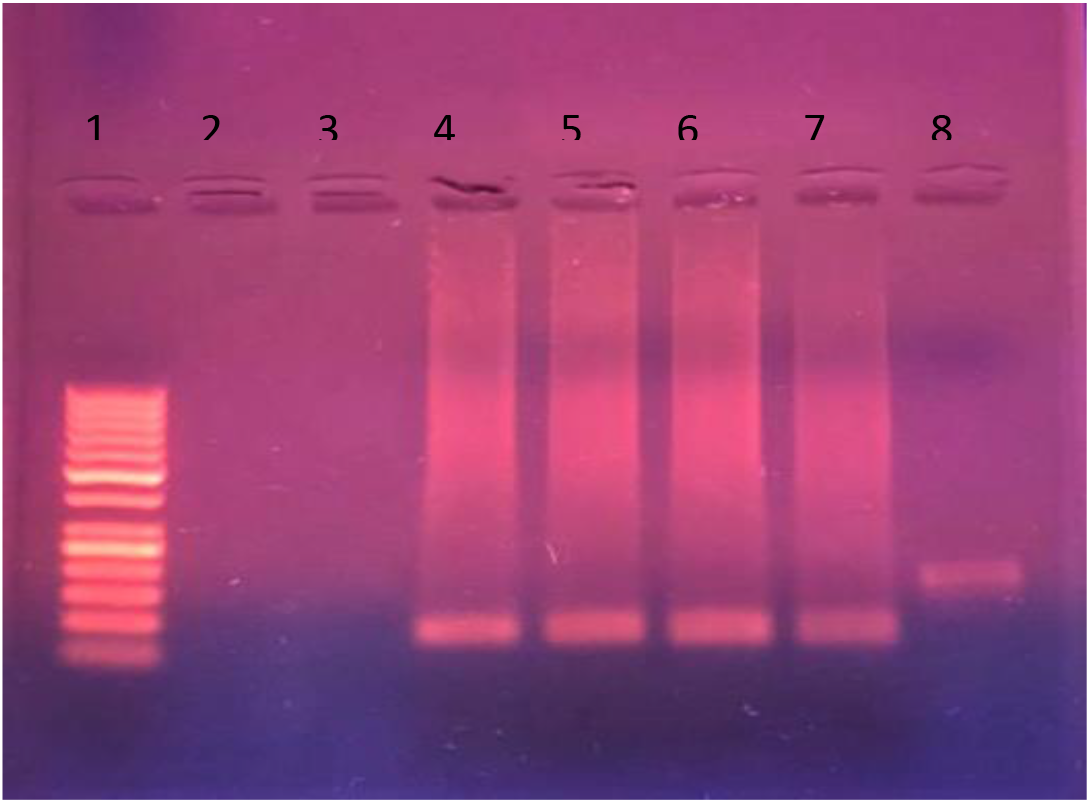
Agarose gel electrophoresis of PCR amplification for EBV detection. Lane 1: DNA marker (50bp), Lane 2-3: Negative, Lane 4-7: EBV-1 (75bp), Lane 8: EBV-2 (168 bp)

**Figure 2a.**
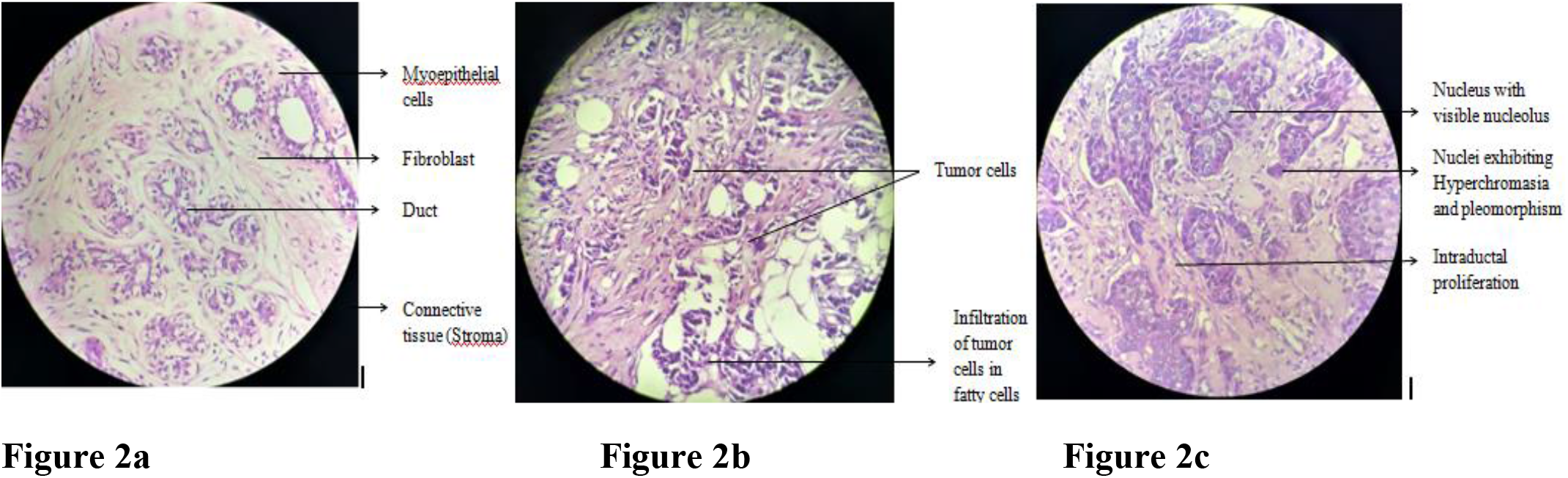
Normal Breast Tissue (control group) (40 X), **Figure 2b** Invasive ductal carcinoma (EBV –ive) (40 X), **Figure 2c** Invasive ductal carcinoma (EBV +ive) (40 X).

### 3.3. Frequency of EBV and EBV genotypes

EBV DNA was detected in 68.3% (41/60) breast cancer patients (Cl= 55.8% – 78.7%), while EBV was not reported in the control group, reported statistically significant (p = 0.001). Of 41 EBV positive patients, 40 (66.66%, 54.1% – 77.3%) patients having EBV-1 while only 1 (1.66%, 0.3% – 8.9%) patient was positive for EBV-2 infection (p < 0.001). The mean (SD) age of EBV DNA-positive and negative patients was 47.098± 12.591 and 49.263± 11.08, respectively, which showed no significant association (p = 0.5, Cl=55.8% – 78.7%). Figure-3 showed the association of EBV genotype with different variables.

**Figure 3.**
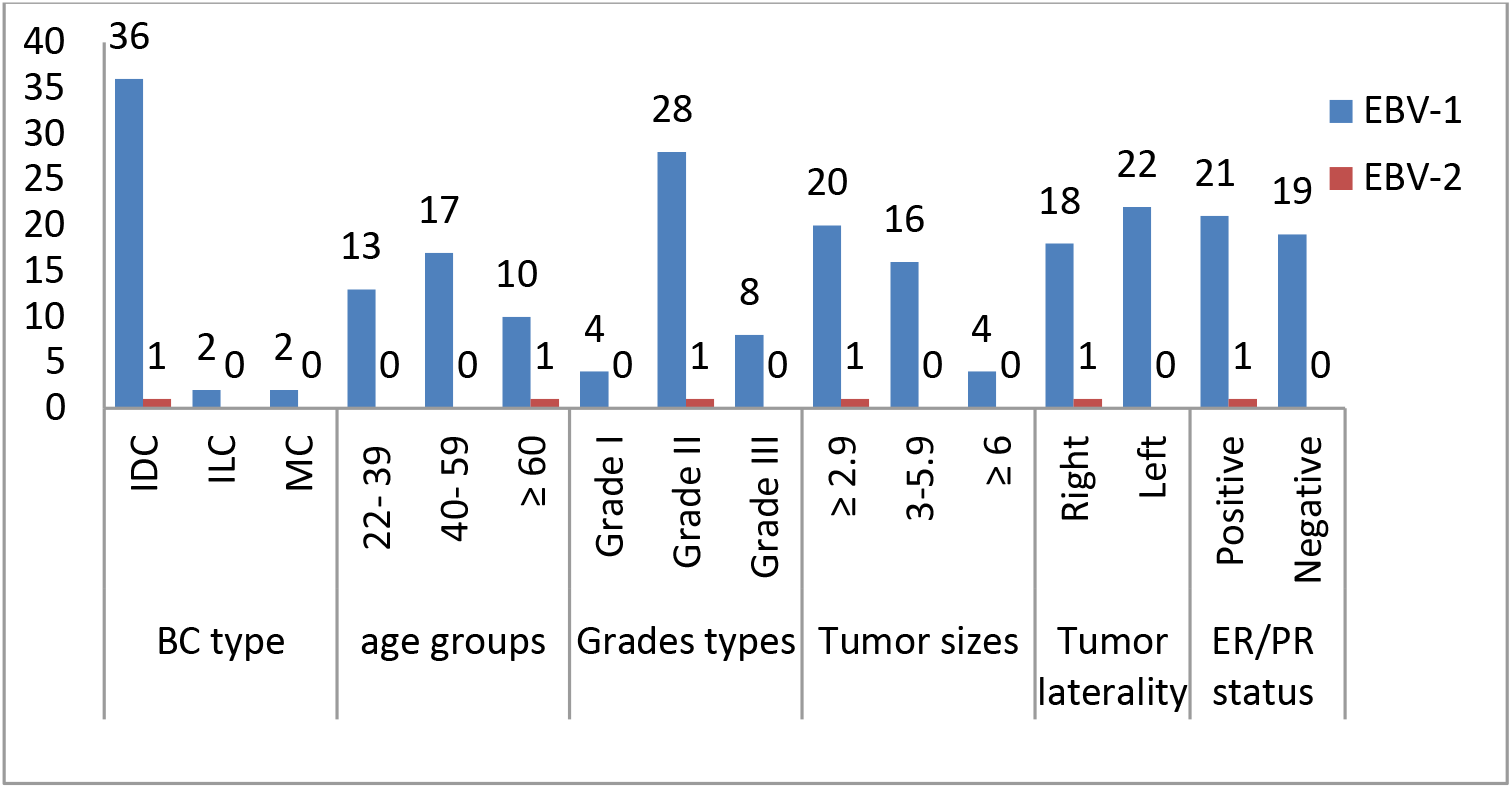
Association of EBV genotype with different variables

### 3.4. Breast cancer type and EBV genotypes

Of 41 positive EBV breast cancer patients, 29.2% (37) were associated with Invasive ductal carcinoma, 48.7% were related with ILC, and 48.7% with metaplastic carcinoma. Of 37 IDC-positive patients, 97.29% (n=36) were positive for EBV-1 and 2.7% (n=1) for EBV-2, while all positive samples of ILC and metaplastic carcinoma were found positive for the EBV-1 genotype. EBV positivity showed no statistically significant association with the histological type (p =0.209).

### 3.5. Age and EBV

The mean (±SD) age of patients with and without EBV were 47.09 (± 12.59) years and 49.26 (± 11.07) years, respectively, showed no statistically significant association (P = 0.476). Table 4 showed EBV frequency in different breast cancer types according to age groups. EBV detection in the age group ≥60 years was higher than the other groups. The highest (78.5%) frequency of EBV-positive samples was reported in IDC patients aged ≥60 years. No statistically significant association was observed between age and frequency of EBV positivity (p= 0.1055).

**Table 4.** EBV frequency in different BC types according to age groups

| AGE | BC TYPE |  |  |  |  |  |  |  |  |  |  |  | Total |  |  | p-value |
| --- | --- | --- | --- | --- | --- | --- | --- | --- | --- | --- | --- | --- | --- | --- | --- | --- |
|  | IDC |  |  | MC |  |  | ILC |  |  | SC |  |  |  |  |  |  |
|  | EBV |  | Total | EBV |  | Total | EBV |  | Total | EBV |  | Total | EBV |  | Total |  |
|  | +ve | -ve |  | +ve | -ve |  | +ve | -ve |  | +ve | -ve |  |  |  |  |  |
| 22-39 | 11 | 3 | 14 | 1 | - | 1 | 1 | - | 1 |  |  |  | 13 | 3 | 16 | 0.768 |
| 40-59 | 15 | 11 | 26 | 1 | - | 1 | 1 | - | 1 | - | 1 | 1 | 17 | 12 | 29 | 0.417 |
| ≥ 60 | 11 | 3 | 14 | - | - | - | - | 1 | 1 | - | - | - | 11 | 4 | 15 | 0.86 |
| Total | 37 | 17 | 54 | 2 | - | 2 | 2 | 1 | 3 | - | 1 | 1 | 41 | 19 | 60 | 0.378 |

### 3.6. Tumor grade and EBV genotypes

Of EBV-positive patients, 4 (9.7 %) have tumor grade I, tumor grade II was found in 28 (68.2 %), and tumor grade III was found in 9 (21.9 %) patients (p= 0.4811) (table 5). Only 1 patient of tumor grade II was EBV-2 positive. The highest ratio of IDC breast cancer (91.6%) was reported in patients with tumor grade III and in patients > 60 years. Table 4 showed EBV frequencies in different age groups in breast cancer types and tumor grades. No statistically significant association was reported between EBV positivity and tumor stage (P = 0.356). The mean age of EBV breast cancer patients of grade I, grade II, and grade III were 40.25 (± 7.14), 50.37 (±13.12), and 41.11 (± 9.31) years old, respectively, and statistically showed significant association (p=0.095).

**Table 5.**
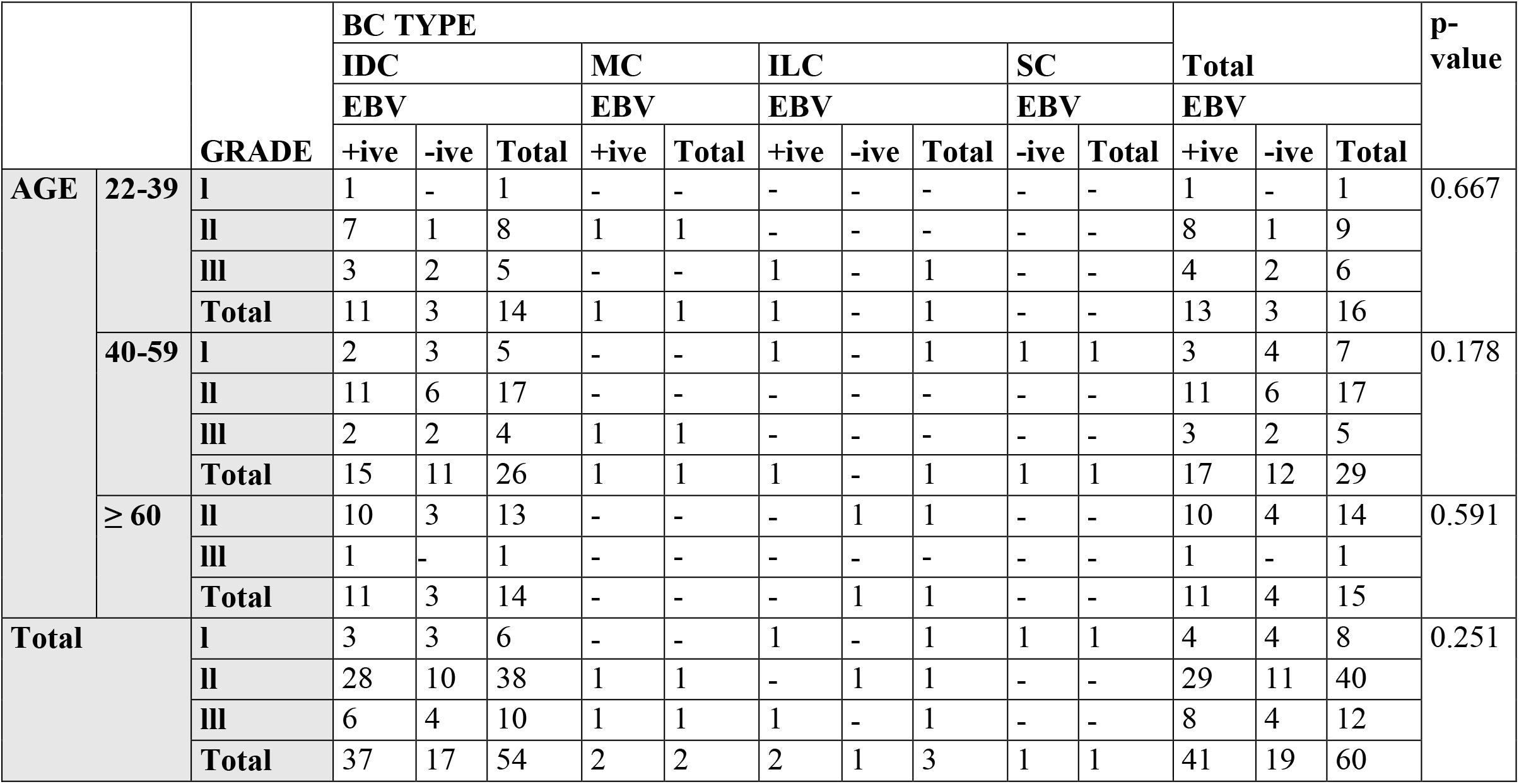
EBV frequency in different age groups in BC types and tumor grades

**Table 6.**
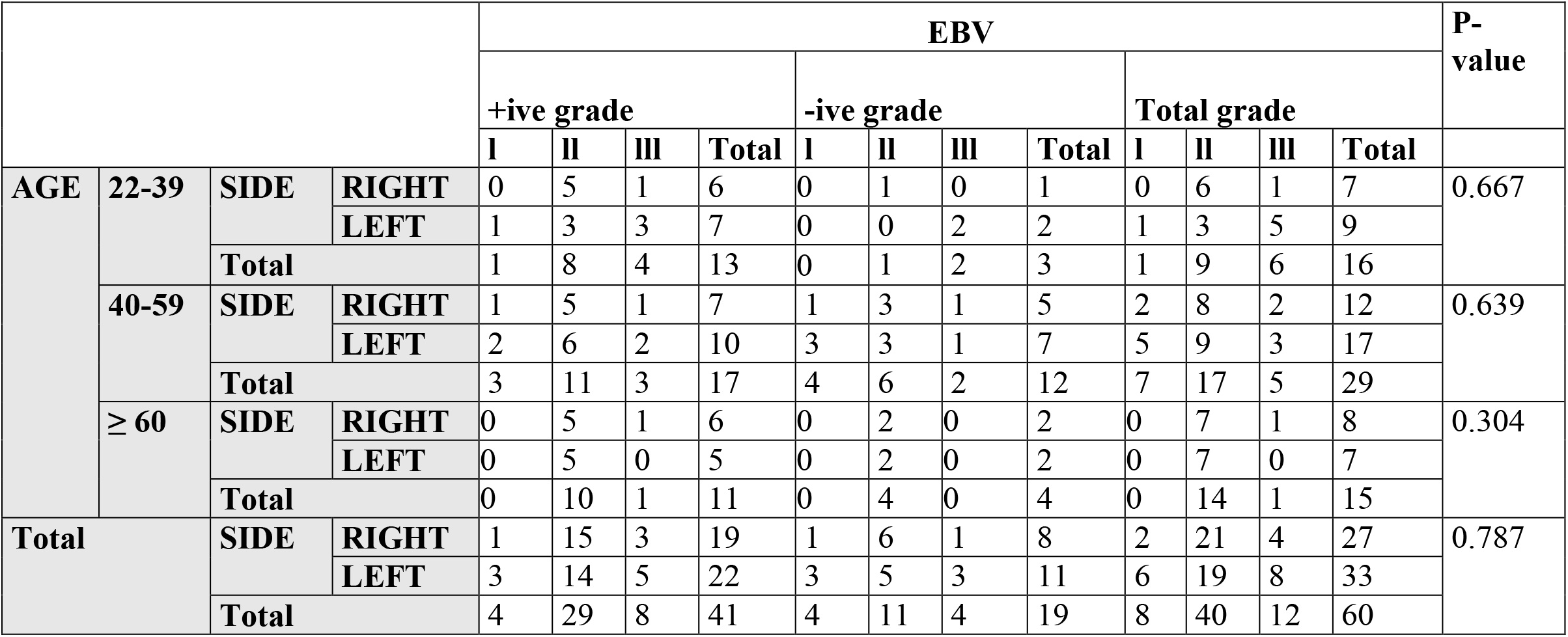
Frequency of EBV according to tumor grade and size **(cm)** in different age groups (years)

### 3.7. Tumor size and EBV genotypes

The tumor size ranged from 0.1-8.5cm in diameter in EBV-1 positive breast cancer patients, 2.2cm in EBV-2 positive patients, and 0.6cm - 6.1cm in EBV negative breast cancer patients. Nineteen (95 %) EBV-1 and 1 (5 %) EBV-2 positive patients have tumor size of 0.1-2.9cm, 17 (41.4 %) EBV-1 patients belonged to tumor size 3-5.9cm and 4 (9.7 %) to tumor size ≥6 (table 4). No statistically significant association was observed between tumor sizes and frequency of EBV (p= 0.976).The samples of tumor grade II having tumor size 0.1-2.9cm showed the highest frequency of EBV in the age group of ≥60 (58.3%), followed by tumor grade II and tumor size 0.1-2.9cm in the age group of 22-39 years ( 38.4%).

### 3.8. Tumor laterality and EBV genotypes

On basis of tumor laterality, 17 (41.4%, Cl= *44.2% – 78.5%*) EBV positive samples were associated with right breast tumor whereas 24 (58.5%, Cl= *55.8% – 84.9%*) were related to left breast tumor. EBV-2 positive samples were associated with the right breast tumor. Regarding the association with ER, tumor laterality showed no statistically significant association (p= 0.533).

### 3.9. ER/PR status and EBV

ER/PR status was positive in 22 (53.6%) EBV-positive patients, whereas 19 (46.34%) were negative for ER/PR status, showed no significant association (p=0.773). An EBV-2 positive patient was also positive to ER/PR status. It was analyzed that the highest ratio of 38.4% of ER/PR receptor non-availability status was in left breast cancer tumors in the age group ≥60 years, followed by left breast tumor samples among the age group 22-39 years and right breast tumor samples among age group> 60 years. Figure-4 showed EBV frequency in tumor laterality and ER/PR status according to age groups.

**Figure 4.**
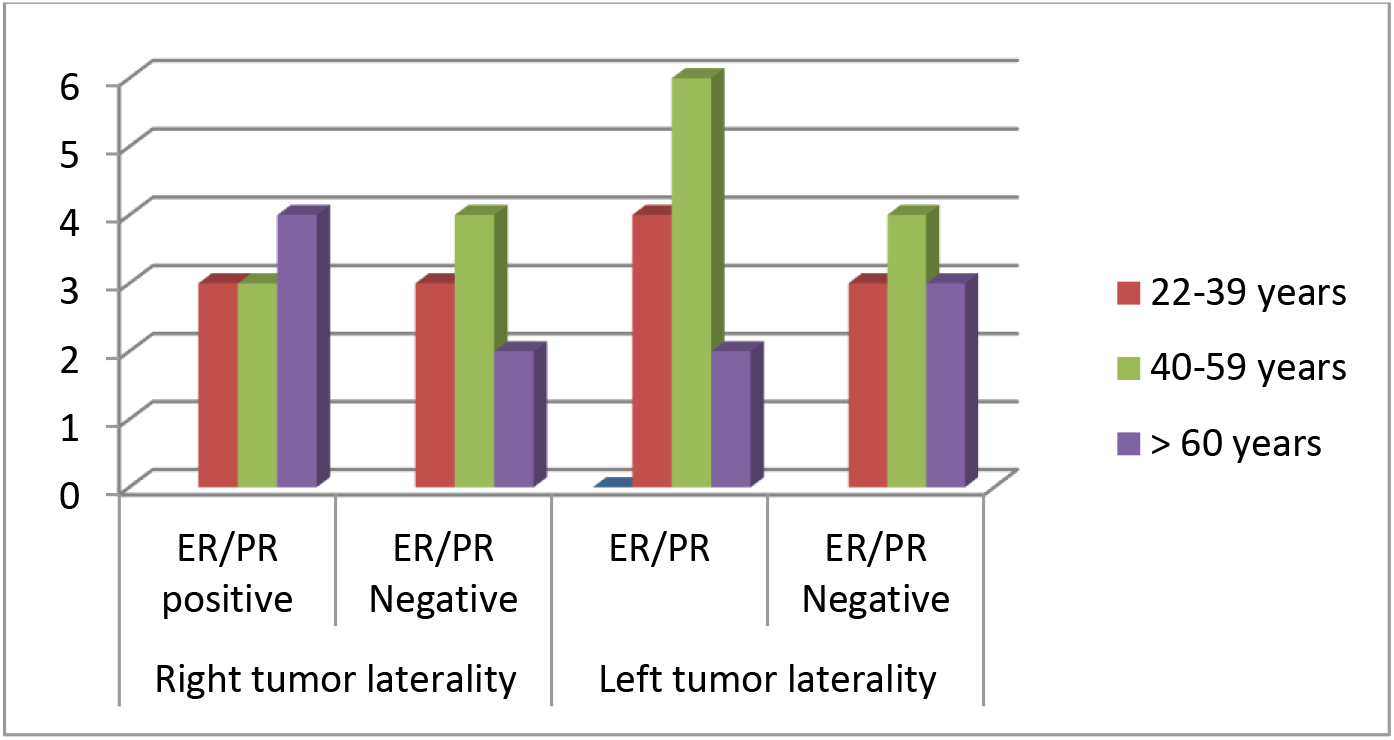
EBV percentage in tumor laterality and ER/PR status according to age groups

## 4. Discussion

Breast cancer – a most frequent malignant tumor in women (Pradhan et al., 2017) is a multifactorial disease that results from the interaction of several predisposing factors (Schwingel et al., 2019), including infectious agents, especially viruses. Despite various other factors, possible viral etiology for breast cancer prompted the researchers to propose few viruses like EBV responsible for its progression. In breast tissue and milk ducts, EBV has been found to infect mammary epithelial cells (Joshi *et al*., 2009).

PCR is potentially a highly sensitive and specific method to detect EBV DNA (). In this study, the use of PCR confirmed the presence of EBV in breast cancer tissue and was comparable with previous PCR results (Sharifpour *et al*., 2019; Naushad *et al*., 2017; Fawzy *et al*., 2008, Arbach et al., 2006, HassabEl-Naby et al., 2017). EBV was localized to infiltrated lymphocytes rather than malignant breast cells (Khan et al., 2011), Therefore, to depend solely on PCR based studies to determine the relationship between EBV DNA and breast cancer risk is inappropriate in infiltrating lymphocytes (Ballard, 2018; Hamad et al., 2020). Reports have indicated an EBER negative form of EBV infection could be present in breast tumor cells (Ballard, 2018). The prevalence rate was highest when PCR targeted the EBER and the reiterated *Bam*H1W sequence. Thus a number of investigators used immunohistochemistry as either the primary or secondary detection system for confirmation, allowing identification of cell type (differentiating infected tumor cells from infiltrating lymphocytes) (Sally et al., 2004). In this study, 60 patients were analyzed for the presence of EBV, of which 68.3% of breast cancer patients were positive for EBV DNA. Different studies reported EBV detection in breast cancer patients worldwide (Hamad et al., 2020). Various studies reported a higher prevalence of viral infections in breast cancer patients than in the control group (Fozuni et al., 2020). In the current study, EBV was not detected in the control group while 66.66% patients were having EBV-1 while only 1.66% patient was positive for EBV-2 infection. Salahuddin et al., in 2018, established that EBV-1 was the predominant genotype in all kinds of lymphoma populations from different hospitals of Islamabad, Pakistan. The highest number of patients infected with EBV-1 and showed a statistically significant association. Joshi et al., 2009 also found an alliance between EBNA −1 gene expression in tumor cells and breast cancer showing 55 % positive results. In our neighboring countries, the predominant genotype was EBV-1, as reported by Tabibzadeh et al. 2020 in the Iranian population (91.2%) and Janani et al. 2015 in Indians (100%). Salahuddin et al. 2018 also showed that EBV-1 (90.7%) was the most prevalent type in the lymphoma population of Pakistan. It was reported that EBV-1 is found globally, predominantly in American, Chinese, European, and South-East Asian populations, whereas type 2 is predominantly found in Africa (Ayee et al., 2020), Alaska, and Papua New Guinea (Zanella e al., 2019). In this study, on the basis of histological types of breast cancer, Invasive ductal carcinoma was reported as the leading type of breast cancer in females, and the majority of EBV was detected in IDC patients. Sharifpour et al. 2019 reported a strong association between EBV and IDC. Both ductal and lobular breast cancer reported a relatively higher EBV prevalence than mixed carcinomas or other types of breast tumors. These differences in prevalence may be because EBV infection elevates breast cancer risk in some specific types of breast carcinoma (Trabelsi et al., 2008).

This study showed significant results indicated to the presence of EBV in the Invasive ductal carcinoma breast cancer regardless of the grade of differentiation. EBV DNA presence in breast cancer patients can differ between EBV in response to various factors, and the genotype frequency was also observed for the specific risk parameters. In this study, age-stratification analysis, no statistically significant association was observed between age and frequency of EBV positivity (p= 0.1055), supported by a study on the frequency of EBV in the Pakistani lymphoma population (Salahuddin et al., 2018).

El-Sheik e al. 2021 reported the risk for an Egyptian woman developing breast cancer increases with age, the majority in women older than 50 years, mentioning that women after the age of 55 went through menopause and had a higher risk of breast cancer due to prolonged exposure of breast cells to estrogen and progesterone. It was reported that menopausal status was associated with breast cancer risk in Pakistan (Sohail et al., 2013, Shamsi et al., 2013). In Asia, breast cancer incidence peaks among women in their forties, whereas in the United States and Europe, it peaks among women in the sixth decade of their life. In this study, ER/PR status was positive in 22 (53.6%) EBV-positive patients, whereas 19 (46.34%) were negative for ER/PR status, showed no significant association (p=0.773). Khan et al. 2011 reported 47.5% of the EBV-positive cases but found no correlation between EBV infected infiltrating lymphocytes and ER-negative breast tumors. Other investigators reported that ER negativity appears to be the dominant factor in the association between EBV and the aggressive breast cancer subtypes. Concerning the odds ratio, EBV infection is 3.57 times more likely in ER-negative tumors (P <0.001), mentioning the higher proportion of ER-negative cells in the basal layer makes these cells more susceptible to the transfer of EBV infection from infiltrating lymphocytes (Ballard et al., 2018). Multiple factors attribute variation in peak incidence age, including geographic variation, racial/ethnic background, genetic variation, lifestyle, environmental factors, socioeconomic status, utilization of screening mammography, stage of disease at diagnosis, and the availability of appropriate care (Leong et al., 2010). HassabEl-Naby et al. 2017 reported that epidemiological variation in EBV infections might be due to variance in age at the time of acquiring primary EBV infection and diversity in the methodologies used for detecting the virus and different EBV-derived proteins or nucleic acids investigated. The inconsistency could be attributed to several aspects: a study design that involved a specific histological type of breast cancer, differences in methodologies or techniques (PCR, ISH, and IHC), dissimilarities in the archival materials (paraffin-embedded and fresh/frozen tissues) (Glaser et al., 2017), geographical and immunological differences and ethnic origin as stated by Glaser et al., (2017) and Thompson (2004).

Regarding the association of EBV presence with tumor grade, 9.7 % have tumor grade I, tumor grade II was found in 8.2 %, and tumor grade III was found in 21.9 % of patients. It was supported by Sharifpour et al. 2019 that a high prevalence of EBV DNA (58.33%) was reported in grade III. However, this study showed no significant association. Zekri et al., 2012, also reported no statistically significant difference between EBV presence and tumor grade in Egyptian (*p* = 0.860) and (Iraqi *p* = 0.976) women. However, Mazouni et al. 2011 reported a significant increase of EBV-positive samples with an increase in tumor grade, with the rate of 16.2% for grade I, to 32.0% for grades II and 46.4% for grades III. The variation of the study might be due to the large sample (n=196). In this study, most EBV breast cancer cases were associated with the left breast tumor based on tumor laterality. This reason might be that the left breast is bulkier, and the upper outer quadrant has a relatively larger volume of breast tissue than the right breast (Kulkarni et al., 2012). The study was also conducted to analyze the EBV frequency based on the laterality ratio, which indicated that 58.5% of tumors positive for EBV were left breast tumors. According to Sughrue and Brody, 2014, Breast cancer was more diagnosed in the left than right breast, regardless of other factors, and established a significant association in breast cancer laterality ratio and increasing age. Dane et al. 2001 also suggested more eminent hypersensitivity on the left side than the right part of the body.

In recent studies, about 19 (31.6%) breast samples showed negative for EBV DNA. Some other viruses like mouse mammary tumor virus (MMTV), Human papillomavirus (HPV), Cytomegalovirus (CMV) have been inspected (Naushad *et al*., 2017; Richardson *et al*., 2015; Glenn *et al*., 2012). The function of the viruses listed needs further examination. The screening of EBV DNA for women with breast cancer should be carried out with the above data to manage and regulate the spread of EBV infection.

## 5. Conclusion

The current research indicates breast carcinoma patients with a high prevalence of 68.3% EBV DNA type 1 observed in formalin-fixed paraffin-embedded tissue. The findings of this study indicate that in Khyber Pakhtunkhwa, Pakistan, EBV had a significant role in invasive ductal breast cancer of tumor grade III among age group III. Approximately 31.6% of the breast samples were negative for EBV DNA. Also, the microscopic analysis of slides was conducted, showing that the virus might induce cancer but haven’t had any effect on cancer histology. Given the prevalence of breast cancer, the function of EBV in the cause or progression of breast cancer, even in a limited subset of cases, will be critical. In future studies, the connection of other viruses, including Human Cytomegalovirus, Papilloma Viruses, and Mammary Mouse viruses, in breast cancer progression should be investigated. Also, it is needed to determine the precise function of EBV in the etiology or advancement of breast cancer, as this will aid in the development of prevention, early detection, and treatment strategies.

## Ethical approval

This article does not contain any studies with human participants performed by any of the authors.

## Authors’ contributions

Yusra Ilyas devised the research goals and objectives, conducted experiments. Sanaullah Khan and Yusra Ilyas analyzed the results. Naveed Khan provided the resources and managed data curation. Sanaullah Khan and Sobia Attaullah reviewed the manuscript. The final draft of this work was read and acknowledged by all authors.

## References

Ahmad, A., 2019. Breast cancer statistics: recent trends. Breast Cancer Metastasis and Drug Resistance, 1–7. https://doi.org/10.1007/978-3-030-20301-6_1.

Al Hamad, M., et al., 2020. Human mammary tumor virus, human papillomavirus, and Epstein-Barr virus infection are associated with sporadic breast cancer metastasis. Breast Cancer:Basic Clin. Res., 14. https://doi.org/10.1177/1178223420976388.

Alibek, K., Kakpenova, A., Mussabekova, A., Sypabekova, M., Karatayeva, N., 2013. Role of viruses in the development of breast cancer. Infect. Agents Cancer, 8(1), 1–6. https://doi.org/10.1186/1750-9378-8-32.

Arbach, H., et al., 2006. Epstein-Barr virus (EBV) genome and expression in breast cancer tissue: effect of EBV infection of breast cancer cells on resistance to paclitaxel (Taxol). J. Virol., 80(2), 845–853. https://doi.org/10.1128/JVI.80.2.845-853.2006.

Asif, H. M., Sultana, S., Akhtar, N., Rehman, J. U., Rehman, R. U., 2014. Prevalence, risk factors and disease knowledge of breast cancer in Pakistan. Asian Pac. J. Cancer Prev., 15(11), 4411–4416. https://doi.org/10.7314/APJCP.2014.15.11.4411.

Ayee, R., Ofori, M. E. O., Wright, E., Quaye, O., 2020. Epstein Barr virus associated lymphomas and epithelia cancers in humans. J. Cancer., 11(7), 1737. https://dx.doi.org/10.7150%2Fjca.37282.

Ballard, A. J., 2018. Epstein - Barr virus is Associated with Aggressive Subtypes of Invasive Ductal Carcinoma of Breast (Her2+/ER-and Triple Negative) and With Nuclear Expression of NF? B p50. J Breast Cancer., 68–75. https://doi.org/10.19187/abc.20185268-75.

Bano, R., Ismail, M., Nadeem, A., Khan, M., Rashid, H., 2016. Potential Risk Factors for Breast Cancer in Pakistani Women. Asian Pac. J. Cancer Prev, 17(9), 4307–4312.

Böhm D., 2010. Investigation of the mechanism of Epstein-Barr virus latent membrane protein 1-mediated NF-κB activationty. The Free University of Berlin, Diss. http://dx.doi.org/10.17169/refubium-15145.

Dane, Ş., Erdem, T., Gümüştekin, K., 2001. Cell-mediated immune hypersensitivity is stronger in the left side of the body than the right in healthy young subjects. Percept. Mot. Ski., 93(2), 329–332. https://doi.org/10.2466%2Fpms.2001.93.2.329.

Dumitrescu, R. G., Cotarla, I., 2005. Understanding breast cancer risk - where do we stand in 2005? J. Cell. Mol. Med, 9(1), 208–221. https://doi.org/10.1111/j.1582-4934.2005.tb00350.x.

El-Naby, N. E. H., Mohamed, H. H., Goda, A. M., Mohamed, A. E. S., 2017. Epstein-Barr virus infection and breast invasive ductal carcinoma in Egyptian women: a single center experience. J Egypt Natl Canc Inst., 29(2), 77–82. https://doi.org/10.1016/j.jnci.2017.02.002

El-Sheikh, N., et al., 2021. Assessment of human papillomavirus infection and risk factors in Egyptian women with breast cancer. Breast Cancer:Basic Clin. Res., 15. https://doi.org/10.1177%2F1178223421996279.

Farrell, P. J., 2019. Epstein–Barr Virus and Cancer. Annu. Rev. Pathol, 14(1), 29–53.

Fawzy, S., Sallam, M., Awad, N. M., 2008. Detection of Epstein–Barr virus in breast carcinoma in Egyptian women. Clin. Biochem, 41(7-8), 486–492. https://doi.org/10.1146/annurev-pathmechdis-012418-013023.

Fessahaye, G., Elhassan, A. M., Elamin, E. M., Adam, A., Ghebremedhin, A., Ibrahim, M. E., 2017. Association of Epstein - Barr virus and breast cancer in Eritrea. Infect. Agents Cancer, 12, 62. https://doi.org/10.1186/s13027-017-0173-2.

Fozuni, E., Mollaei, H. R., Iranpour, M., Afshar, R. M., 2020. Evaluation frequency of Human Herpes Virus type 8 in Patients with Breast Cancer. https://doi.org/10.21203/rs.3.rs-60897/v1

Gannon, O. M., Antonsson, A., Bennett, I. C., Saunders, N. A., 2018. Viral infections and breast cancer – A current perspective. Cancer Lett., 420, 182–189. https://doi.org/10.1016/j.canlet.2018.01.076.

Ghoncheh, M., Mohammadian-Hafshejani, A., Salehiniya, H., 2015. Incidence and mortality of breast cancer and their relationship to development in Asia. Asian Pac. J. Cancer Prev., 16(14), 6081–6087.

Glaser, S. L., Canchola, A. J., Keegan, T. H., Clarke, C. A., Longacre, T. A., Gulley, M. L., 2017. Variation in risk and outcomes of Epstein–Barr virus-associated breast cancer by epidemiologic characteristics and virus detection strategies: an exploratory study. Cancer Causes & Control, 28(4), 273–287. https://doi.org/10.1007/s10552-017-0865-3

Glaser, S. L., Hsu, J. L., Gulley, M. L., 2004. Epstein-Barr virus and breast cancer: state of the evidence for viral carcinogenesis. Cancer Epidemiol. Biomark. Prev., 13(5), 688–697.

Glenn, W. K., Heng, B., Delprado, W., Iacopetta, B., Whitaker, N. J., Lawson, J. S., 2012. Epstein-Barr virus, human papillomavirus and mouse mammary tumour virus as multiple viruses in breast cancer. PloS one, 7(11), e48788.

Gulley, M. L., 2001. Molecular diagnosis of Epstein-Barr virus-related diseases. J Mol Diagn., 3(1), 1–10. https://doi.org/10.1016/S1525-1578(10)60642-3.

Harbeck, N., et al., 2019. Breast cancer (Primer). Nat. Rev. Dis. Primers, 5(1):66. http://doi:10.1038/s41572-019-0111-2.

Hippocrate, A., Oussaief, L., Joab, I., 2011. Possible role of EBV in breast cancer and other unusually EBV-associated cancers. Cancer Lett, 305(2), 144–149. https://doi.org/10.1016/j.canlet.2010.11.007.

Hu, H., et al., 2016. Epstein-Barr Virus Infection of Mammary Epithelial Cells Promotes Malignant Transformation. EBioMedicine, 9, 148–160. https://doi.org/10.1016/j.ebiom.2016.05.025

Huo, Q., Zhang, N., Yang, Q., 2016. Epstein-Barr virus infection and sporadic breast cancer risk: a meta-analysis. PloS one, 7(2), e31656.

Janani, M. K., Malathi, J., Appaswamy, A., Singha, N. R., Madhavan, H. N., 2015. A hospital based pilot study on Epstein-Barr virus in suspected infectious mononucleosis pediatric patients in India. J. Infect. Dev. Ctries., 9(10), 1133–1138. https://doi.org/10.3855/jidc.6199.

Jean-Pierre, V., Lupo, J., Buisson, M., Morand, P., Germi, R., 2021. Main targets of interest for the development of a prophylactic or therapeutic Epstein - Barr virus vaccine. Front. microbiol., 12. https://dx.doi.org/10.3389%2Ffmicb.2021.701611.

Johannsen, E. C., Kaye, K. M., 2015. Epstein-Barr virus (infectious mononucleosis, Epstein-Barr virus-associated malignant diseases, and other diseases. Infect Dis Clin Pract, 7(1602), 1–16.

Joshi, D., Quadri, M., Gangane, N., Joshi, R., Gangane, N., 2009. Association of Epstein Barr virus infection (EBV) with breast cancer in rural Indian women. PLoS One, 4(12), e8180.

Khan, G., Philip, P. S., Al Ashari, M., Houcinat, Y., Daoud, S., 2011. Localization of Epstein-Barr virus to infiltrating lymphocytes in breast carcinomas and not malignant cells. Exp. Mol. Pathol., 91(1), 466–470. https://doi.org/10.1016/j.yexmp.2011.04.018.

Kulkarni, S., et al., 2012. Increased expression levels of WAVE3 are associated with the progression and metastasis of triple negative breast cancer. PloS one, e42895.

Lawson, J. S., and Heng, B., 2010. Viruses and breast cancer. Cancers, 2(2), 752–772. https://doi.org/10.3390/cancers2020752.

Lee, S. M., Park, J. H., Park, H. J., 2008. Breast cancer risk factors in Korean women: a literature review. International nursing review, 55(3), 355–359. https://doi.org/10.1111/j.1466-7657.2008.00633.x.

Leong, S. P., et al., 2010. Is breast cancer the same disease in Asian and Western countries?. World J. Surg., 34(10), 2308–2324. https://doi.org/10.1007/s00268-010-0683-1/

Li, H. P., Chao, M., Chen, S. J., Chang, Y. S., 2010. Epstein-barr virus and its oncogenesis. Human oncogenic viruses, 209–267. https://doi.org/10.1142/9789812833471_0005.

Madhav, M. R., et al., 2018. Epidemiologic analysis of breast cancer incidence, prevalence, and mortality in India: Protocol for a systematic review and meta-analyses. Medicine, 97(52). https://dx.doi.org/10.1097%2FMD.0000000000013680.

Majeed, W., Aslam, B., Javed, I., Khaliq, T., Muhammad, F., Ali, A., Raza, A., 2014. Breast cancer: major risk factors and recent developments in treatment. APJCP, 15(8), 3353–3358. https://doi.org/10.7314/APJCP.2014.15.8.3353.

Mazouni, C., et al., 2011. Epstein-Barr virus as a marker of biological aggressiveness in breast cancer. Br. J. Cancer, 104(2), 332–337. https://doi.org/10.1038/sj.bjc.6606048.

McPherson K., Steel C.M., Dixon J.M., 2000. Breast cancer epidemiology, risk factors, and genetics. Bmj., 321, 624–628. https://doi.org/10.1136/bmj.321.7261.624.

Menhas, R., Umer, S., 2015. Breast cancer among Pakistani women. Iran. J. Public Health, 44(4), 586–7.

Naushad, W., Surriya, O., Sadia, H., 2017. Prevalence of EBV, HPV and MMTV in Pakistani breast cancer patients: A possible etiological role of viruses in breast cancer. Infect. Genet. Evol., 54, 230–237. https://doi.org/10.1016/j.meegid.2017.07.010.

Pradhan, A., Paudyal, P., Sinha, A. K., Agrawal, C. S., 2017. Grading, staging and Nottingham prognostic index scoring of breast carcinoma. J. pathol. Nepal, 7(1), 1078–1083. https://doi.org/10.3126/jpn.v7i1.16951.

Richardson, A.K., et al., 2015. Cytomegalovirus and Epstein-Barr virus in breast cancer. PloS one, 10(2), e0118989.

Salahuddin, S., et al., 2018. Prevalence of Epstein–Barr Virus Genotypes in Pakistani Lymphoma Patients. Asian Pac. J. Cancer Prev, 19(11), 3153–3159. https://dx.doi.org/10.31557%2FAPJCP.2018.19.11.3153.

Schwingel, D., Andreolla, A. P., Erpen, L. M., Frandoloso, R., Kreutz, L. C., 2019. Bovine leukemia virus DNA associated with breast cancer in women from South Brazil. Sci. Rep., 9(1), 1–7. https://doi.org/10.1038/s41598-019-39834-7.

Shamsi, U., Khan, S., Usman, S., Soomro, S., Azam, I., 2013. A multicenter matched case control study of breast cancer risk factors among women in Karachi, Pakistan. APJCP, 14(1), 183–188. https://doi.org/10.7314/APJCP.2013.14.1.183.

Sharifipour, S., Rad, K. D., 2020. Seroprevalence of Epstein–Barr virus among children and adults in Tehran, Iran. New Microbes New Infect, 34. https://doi.org/10.1016/j.nmni.2019.100641.

Sharifpour, C., et al., 2019. Frequency of Epstein–Barr Virus DNA in Formalin-Fixed Paraffin-Embedded Tissue of Patients with Ductal Breast Carcinoma. Asian Pac. J. Cancer Prev, 20(3), 687–692. https://dx.doi.org/10.31557%2FAPJCP.2019.20.3.687.

Smatti, M. K., Al-Sadeq, D. W., Ali, N. H., Pintus, G., Abou-Saleh, H., Nasrallah, G. K., 2018. Epstein–Barr Virus Epidemiology, Serology, and Genetic Variability of LMP-1 Oncogene Among Healthy Population: An Update. Front. Oncol., 8. https://doi.org/10.3389/fonc.2018.00211

Sohail, A., et al., 2013. Effects of glutathione-S-transferase polymorphisms on the risk of breast cancer: A population-based case–control study in Pakistan. Environ. Toxicol. Pharmacol., 35(2), 143–153. https://doi.org/10.1016/j.etap.2012.11.014.

Sughrue, T., Brody, J. P., 2014. Breast tumor laterality in the United States depends upon the country of birth, but not race. PloS one, 9(8), e103313.

Tabibzadeh, A., et al., 2020. Molecular epidemiology of epstein-barr virus (ebv) in patients with hematologic malignancies. APJCP, 21(3). https://dx.doi.org/10.31557%2FAPJCP.2020.21.3.693.

Thompson, M. P., Kurzrock, R., 2004. Epstein-Barr virus and cancer. Clin. Cancer Res., 10(3), 803–821. https://doi.org/10.1158/1078-0432.CCR-0670-3.

Trabelsi A., Rammeh S., Stita W., Mokni M., Mourou A., Korbi, S., 2008. Detection of Epstein-Barr virus in breast cancers with lymphoid stroma. Ann Biol Clin, 66: 59–62. https://doi.org/10.1684/abc.2008.0191.

Zaheer, S., Shah, N., Maqbool, S. A., Soomro, N. M., 2019. Estimates of past and future time trends in age-specific breast cancer incidence among women in Karachi, Pakistan: 2004–2025. BMC public health, 19(1), 1–9. https://doi.org/10.1186/s12889-019-7330-z.

Zanella, L., Riquelme, I., Buchegger, K., Abanto, M., Ili, C., Brebi, P., 2019. A reliable Epstein-Barr virus classification based on phylogenomic and population analyses. Sci. Rep., 9(1), 1–11. https://doi.org/10.1038/s41598-019-45986-3.

Zekri, A. R. N., et al., 2012. Epstein-Barr virus and breast cancer: epidemiological and molecular study on Egyptian and Iraqi women. J Egypt Natl Canc Inst., 24(3), 123–131. https://doi.org/10.1016/j.jnci.2012.06.001.

